## Supplementary Figures for "Functional recruitment of dynamin requires multimeric interactions for efficient endocytosis"

**Inventory of Supplementary Data**

Supplementary Figure 1: Rescue of dynamin function with dyn2-GFP mutants does not depend on their level of expression

Supplementary Figure 2: The position of the GFP tag does not affect dynamin recruitment at the plasma membrane.

Supplementary Figure 3: Synthesis and characterization of divalent peptides

Supplementary Figure 4: Workflow of automated analysis of ppH data (SVM for final validation)

Supplementary Figure 5: Characteristics of the whole cell recordings in the different conditions of this study.

Supplementary Figure 6: Effect of MyrD15 peptide on CME monitored with the ppH assay, on β2 adrenergic receptor and β1 adrenergic receptor internalization.

Supplementary Table 1: characteristics of the divalent peptides

Supplementary Table 2: Proteins identified by mass spectrometry after dD15-N pull down

Supplementary Table 3: P values for the statistical tests used in Figures 1 and 5.

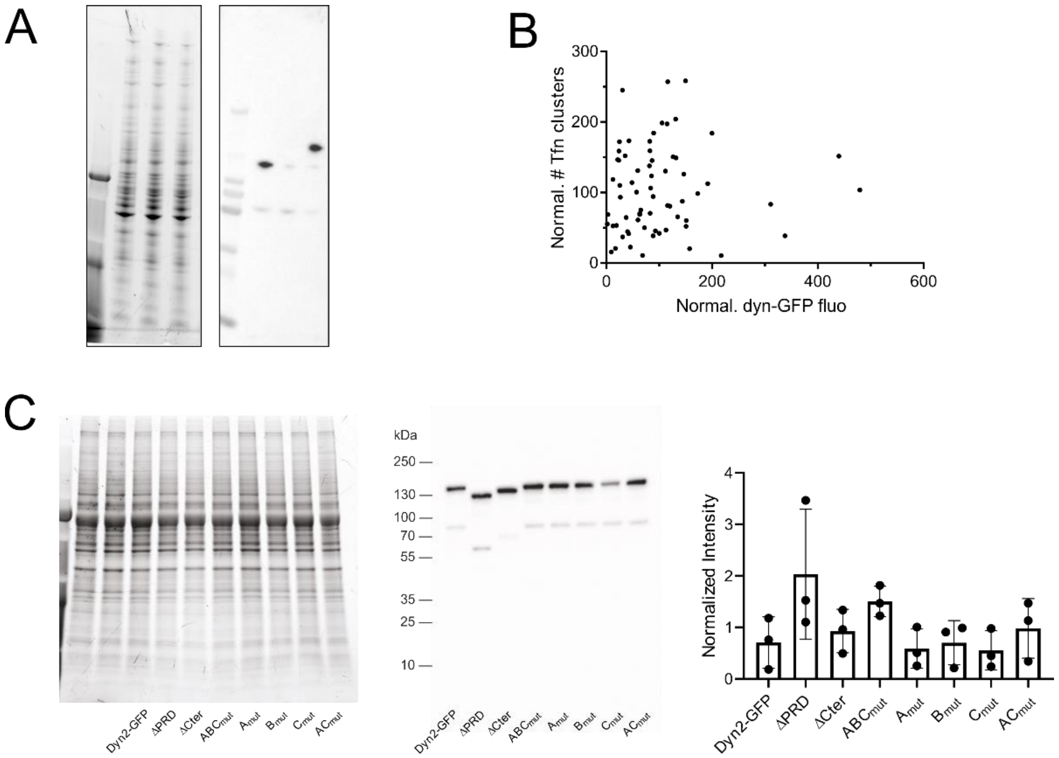

**Supplementary Figure 1: The degree of expression of dynamin mutants does not correlate with internalized transferrin. A**, Full blots corresponding to Figure 1B. Left, stain free detection of total protein. Lane 1, size marker; lanes 2-4, untreated, OHT-treated and OHT-treated + dyn2-GFP-WT transfected lysates respectively. Right, corresponding blot revealed with anti-dynamin antibody. **B**, The fluorescence of each TKO cell transfected with dyn2-GFP-WT was measured as the average in the mask normalized to 100 for each of 4 experimental sessions. There was no significant correlation between the two parameters (Pearson r = 0.058; p = 0.64). **C**, Level of expression of the mutants used in Figure 1D-F estimated by immunoblots using an anti-GFP antibody. Left, stain free detection of total protein. Right, quantification of three independent experiments. The intensities are normalized for each experiment.

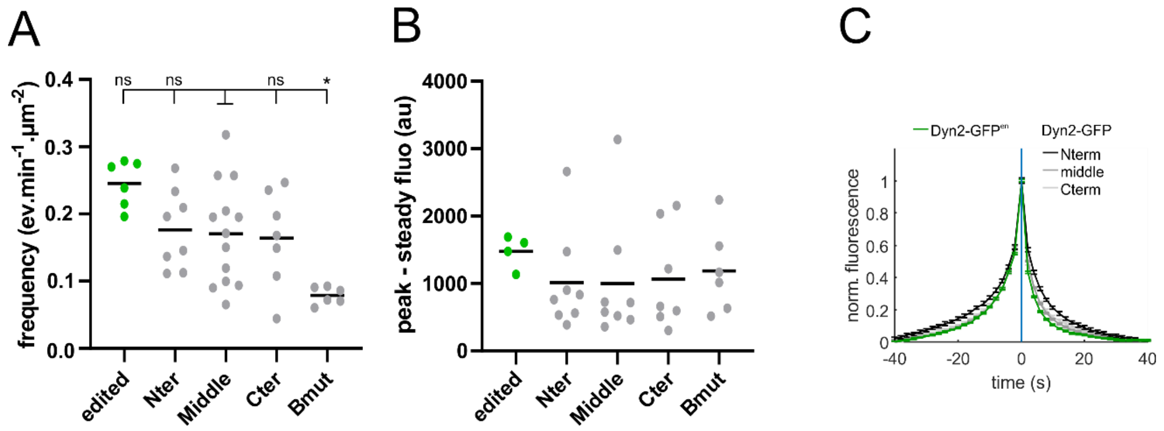

**Supplementary Figure 2: Rescue of dynamin function with dyn2-GFP does not depend on the position of the GFP tag.**

**A**, The frequency of dyn2-GFP recruitment events is the same with different positions of GFP: at the N-terminus, before the PRD and at the C-terminus. Like in Figure 2, the frequency was slightly larger in genome edited dyn2-GFP cells (green bars) but significantly lower with reexpression of dyn2 **B-C**, The amplitudes (B) and kinetics (C) of recruitment are also the same with the three constructs.

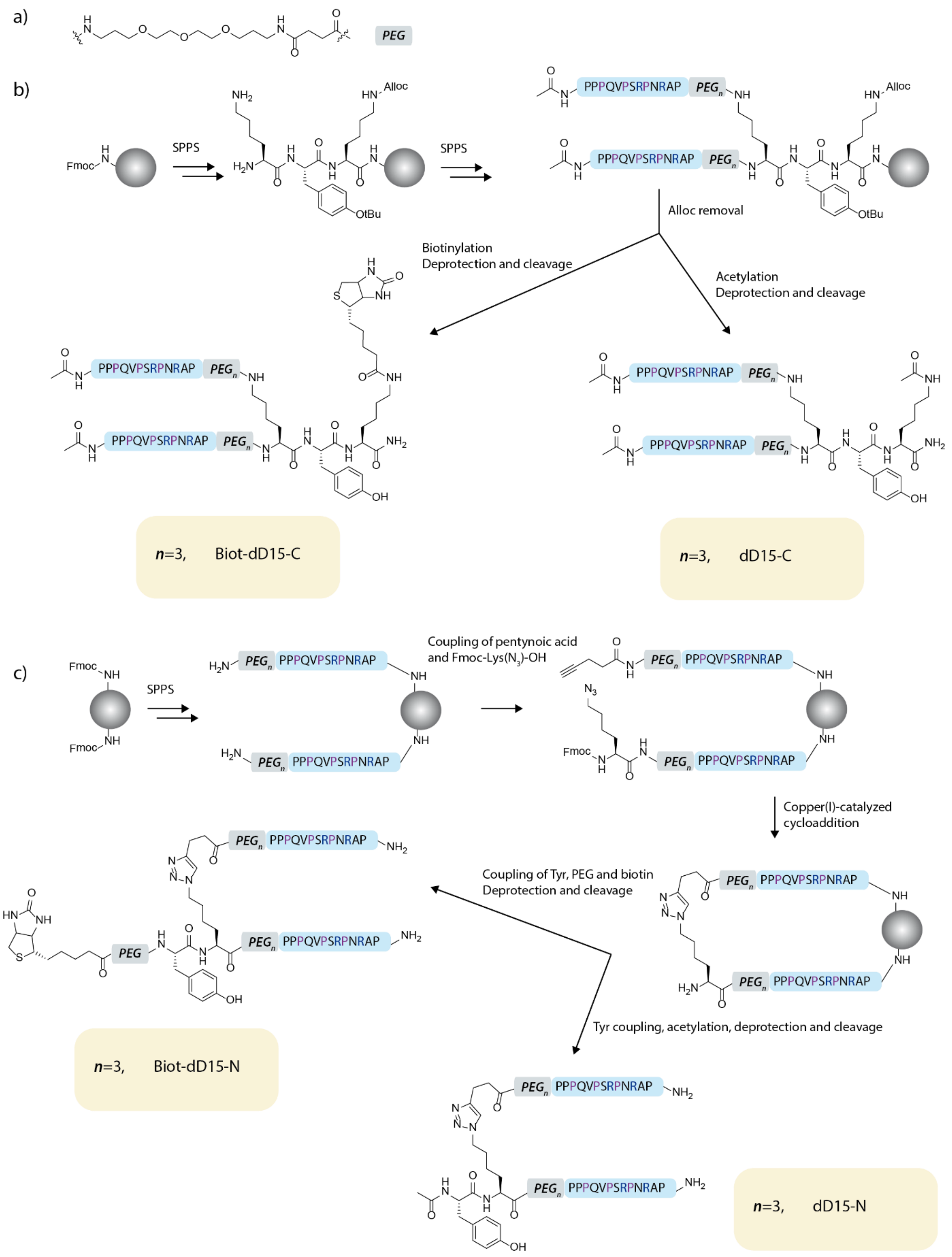

**Supplementary Figure 3**: Synthesis and structure of the divalent peptides. a) Chemical structure of the PEG residue. b) Synthetic schemes and structure of the C-terminally linked divalent peptides. c) Synthetic schemes and structure of the N-terminally linked divalent peptides.

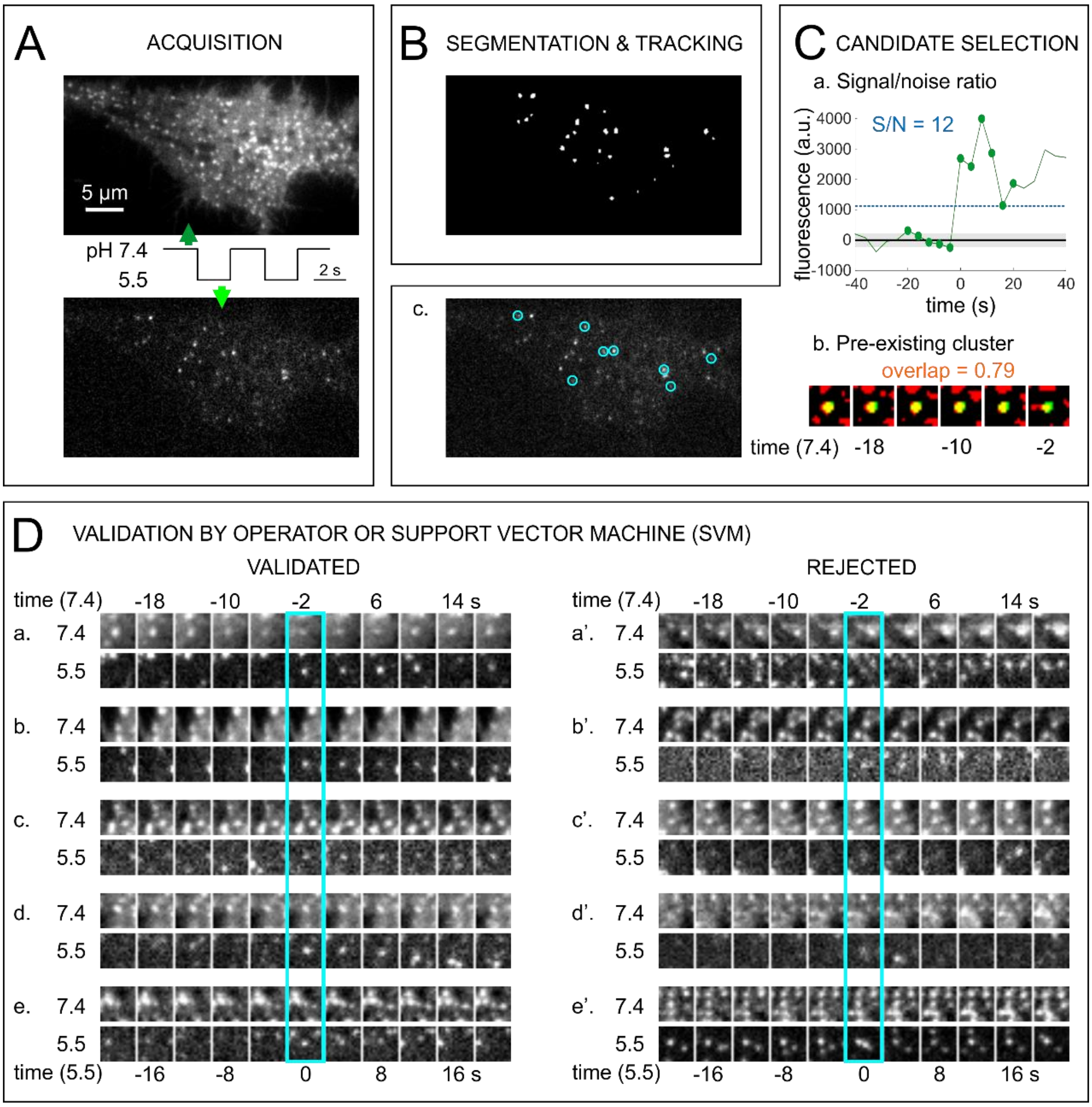

**Supplementary Figure 4**: **Workflow of automated analysis of ppH data.** The four steps (A-D) leading to the characterization of scission events. **A**, Acquisition of images alternatively at pH 7.4 and pH 5.5. Two consecutive images are shown. The image at pH 5.5 is shown with 8x higher brightness than the one at pH 7.4. **B**, Segmentation and tracking of clusters visible at pH 5.5 on the image shown in A. **C**, Selection of candidate events based on two criteria. <u>a. Signal/noise ratio</u>. The average fluorescence in a circle of 2 pixels radius centered on the center of mass of the segmented object is plotted for each frame before and at the start of segmentation (time 0). The noise is estimated as the average ± std before detection (black line, average; gray shading, std). An event is qualified as candidate if the fluorescence at detection divided by the noise estimate (S/N) is bigger than 5. The example (corresponding to event a. in panel D) has S/N = 12.0. <u>b. Pre-existing cluster at pH 7.4.</u> In segmented images, the location of the event (green) is compared to the clusters of TfR visible at pH 7.4 (red) for the five frames preceding the event. The fraction of green pixels overlapping with red (appearing in yellow) gives an estimate of the colocalization of the candidate vesicle with a parent cluster. An event is qualified as candidate when this fraction is greater than 0.2. For the displayed example (event a. in Panel D) the fraction overlap is 0.79. c. The cyan circles mark the clusters segmented in B which passed the criteria. **D**, Final event validation by a human operator or the trained SVM. Left, gallery of events which were validated by a human operator. In all cases (a-e) the acid resistant spot (vesicle) is clearly visible and can be tracked. Right, gallery of events which were selected by the first screen described in C but subsequently rejected by a human operator. In cases a’, d’ and e’ the acid resistant spot cannot be tracked consistently and in cases b’ and c’ it is barely above background. The choices made for validation and rejection, together with 9621 others, have served to train the SVM.

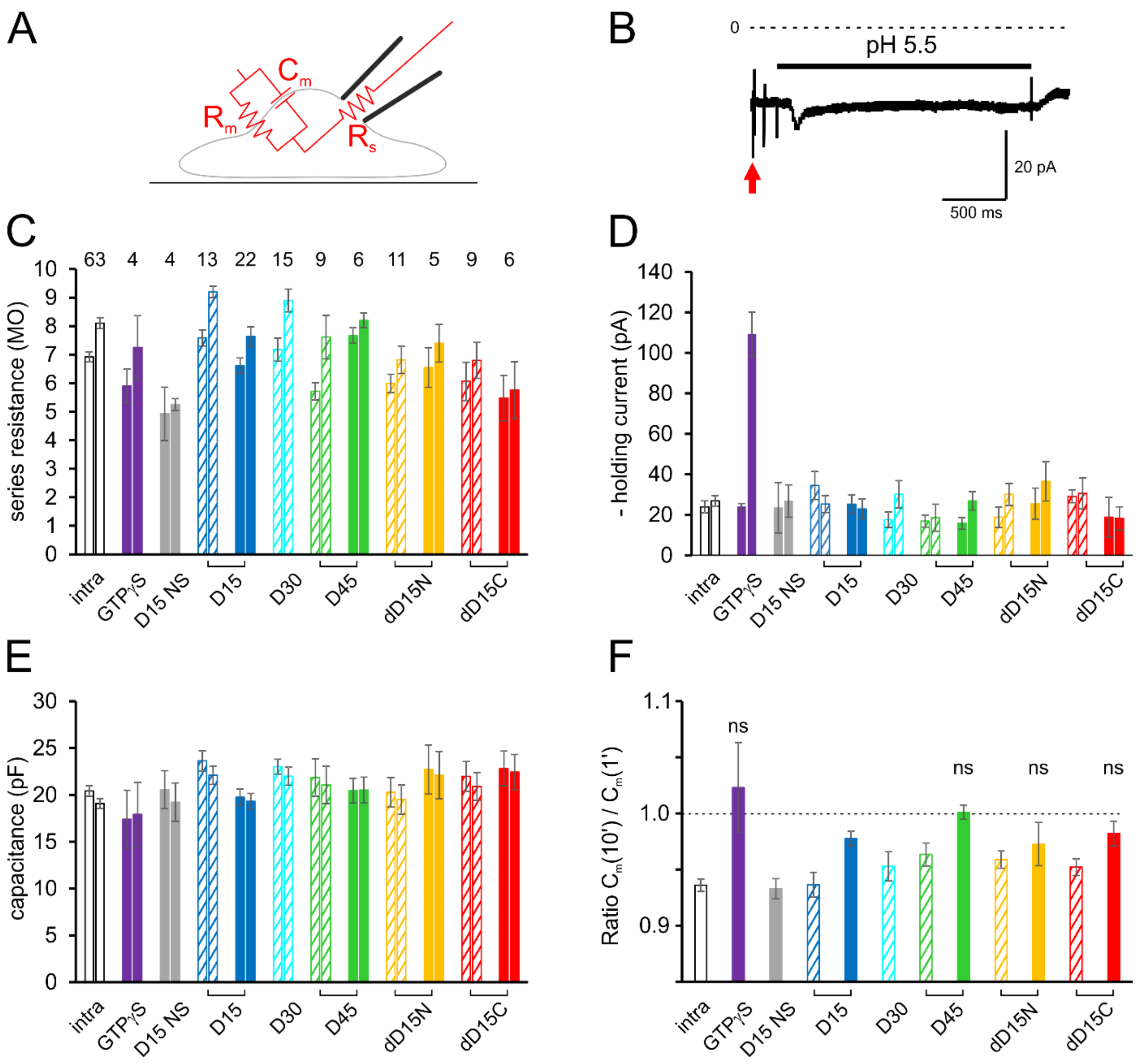

**Supplementary Figure 5: Characteristics of the whole cell recordings in the different conditions of this study. A**, Scheme of the equivalent electrical circuit of a cell recorded in the whole cell configuration. The electrical circuit in *red* is estimated by a voltage pulse (*red arrow* in B) and compensated by the patch clamp amplifier. Series resistance ($R_s$) depends on the pipette resistance and the access to the cell cytoplasm. It reflects the ability of the pipette solution containing the blocking peptides to diffuse into the cell. Membrane capacitance ($C_m$) is proportional to the plasma membrane surface. Membrane resistance ($R_m$) is inversely proportional to the number of open channels. **B**, Example of a current recording of a cell held in voltage clamp at -60 mV. The holding current is -21 pA. The application of solution at pH 5.5 evokes a current of less than 5 pA. Fast transient currents reflect the response to the test voltage step (*red arrow*, -5 mV, 10 ms, see A) and the electrical artefacts of electro-valve openings to exchange the extracellular solutions. **C**, Series resistance $R_s$ for each recording condition (same cells as in Figure 5D), during the 1$^{st}$ minute (left) and the 10$^{th}$ minute (right) of whole cell recording. $R_s$ were maintained below 10 MΩ throughout the recording and do not differ between conditions as compared to the control condition (one way ANOVA and Tukey multiple comparison tests). **D**, Same as C for holding current. There was no significant difference between conditions (one way ANOVA and Tukey's multiple comparison tests)

except for GTPγS at the $10^{th}$ minute where the holding current was much larger ($p < 0.0001$). This is likely due to the activation of GTP dependent trimeric G proteins which activate various channels in the presence of GTPγS, thereby increasing the holding current. **E**, Same as C for membrane capacitance $C_m$. There were no differences between conditions (one way ANOVA and Tukey multiple comparison tests). **F**, Ratios of $C_m$ recorded during the $10^{th}$ minute of recording over $C_m$ recorded during the $1^{st}$ minute. The variation in $C_m$ indicates the balance between exocytosis and endocytosis. In control conditions, the relative membrane capacitance slightly but significant decreases: $C_m(10')/C_m(1') = 0.936 \pm 0.006$ ($p < 10^{-13}$ paired t-test). If endocytosis is blocked, $C_m$ would be expected to stop decreasing or even increase. In four conditions (GTPγS, D44 1 mM, dD15-N 1mM, dD15-C 1 mM) there was no significant decrease in $C_m$ (p = 0.49; 0.67; 0.27; 0.14, respectively). These four conditions correspond to the strongest decrease in endocytic event frequency measured with the ppH assay (see Figure 2D). All graphs indicate average ± SEM.

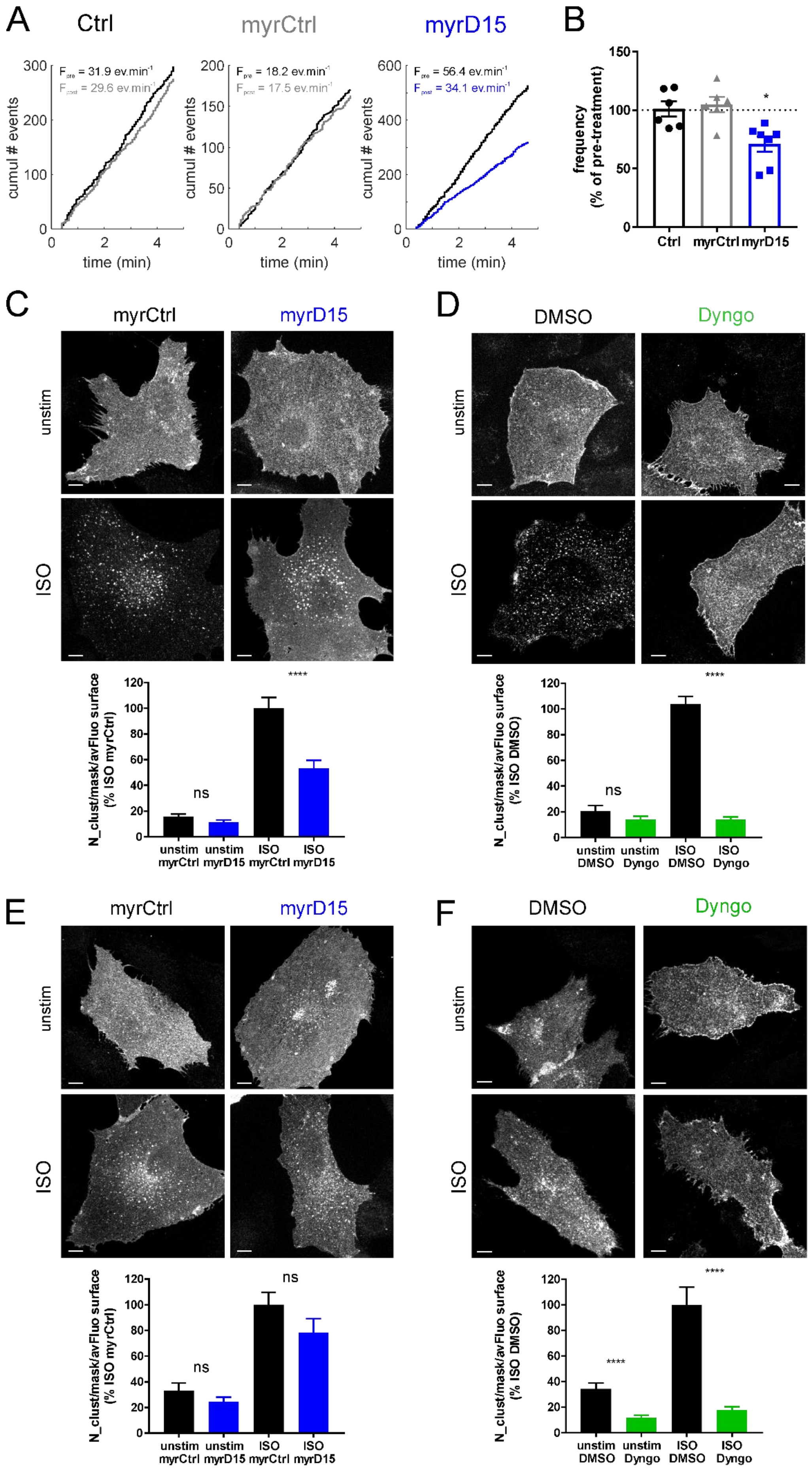
A
Ctrl
myrCtrl
myrD15
cumul # events
time (min)
F_pre = 31.9 ev.min-1
F_post = 29.6 ev.min-1
F_pre = 18.2 ev.min-1
F_post = 17.5 ev.min-1
F_pre = 56.4 ev.min-1
F_post = 34.1 ev.min-1
B
frequency
(% of pre-treatment)
Ctrl
myrCtrl
myrD15
C
myrCtrl
myrD15
unstim
ISO
N_clust/mask/avFluo surface
(% ISO myrCtrl)
ns
****
unstim myrCtrl
unstim myrD15
ISO myrCtrl
ISO myrD15
D
DMSO
Dyngo
unstim
ISO
N_clust/mask/avFluo surface
(% ISO DMSO)
ns
****
unstim DMSO
unstim Dyngo
ISO DMSO
ISO Dyngo
E
myrCtrl
myrD15
unstim
ISO
N_clust/mask/avFluo surface
(% ISO myrCtrl)
ns
ns
unstim myrCtrl
unstim myrD15
ISO myrCtrl
ISO myrD15
F
DMSO
Dyngo
unstim
ISO
N_clust/mask/avFluo surface
(% ISO DMSO)
****
****
unstim DMSO
unstim Dyngo
ISO DMSO
ISO Dyngo

**Supplementary Figure 6: MyrD15 peptide partially blocks CME monitored with the ppH assay, as well as the dynamin-dependent internalization of the CME cargo β2 adrenergic receptor but not of the CIE cargo β1 adrenergic receptor. A**, Examples of ppH assay recordings of 3T3 cells before (PRE, black lines) and after (POST, grey lines) 20 minute incubation with no peptide (Ctrl, left), 10 µM myrCtrl (middle) or 10 µM myrD15 (right)**. B,** Average ± SEM of the frequency of TfR endocytic events measured with the ppH assay in 3T3 cells after 20 minute incubation with no peptide (Ctrl), 10 µM myristoylated control peptide (myrCtrl) or 10 µM myristoylated D15 peptide (myrD15). The dotted line represents the frequency of events before incubation (pre-treatment). **C-F,** Representative spinning disk images (top panels) of BSC-1 cells expressing either β2-adrenergic receptor (C,D) or β1-adrenergic receptor (E,F) and quantification of receptor internalization (expressed as number of clusters per area over average fluorescence at the cell surface) (bottom panels). Cells were either left unstimulated (unstim) or stimulated with 10 µM isoproterenol (ISO). Cells were also pre-treated for 20 minutes with either 10 µM myrCtrl or myrD15 to test the inhibitory effect of myrD15 on internalization of β2- and β1-adrenergic receptors (C,E), or with either DMSO or 30 µM of the dynamin inhibitor Dyngo4a to confirm the dependency of both β2- and β1-adrenergic receptor internalization on dynamin (D, F).

| | Sequence | Formula | Mass (m/z)[a] | | Rt (min), [purity][b] |
|---|---|---|---|---|---|
| | | | Expected M | Obtained $[M+H]^+$ | |
| Ac-D15 | Ac-Y PPPQV PSRPN RAPPG-$CONH_2$ | $C_{80}H_{123}N_{25}O_{21}$ | 1769.9 | 1771.0 | 20.8 [93%] |
| Biot-D15 | Biot-PEG-Y PPPQV PSRPN RAPPG-$CONH_2$ | $C_{102}H_{161}N_{29}O_{27}S$ | 2256.2 | 2257.2 | ND |
| Ac-D44 | Ac-YALGG APPVP SRPGA SPDPF GPPPQ VPSRP NRAPP GVPRI TTSDP-$CONH_2$ | $C_{207}H_{322}N_{60}O_{58}$ | 4576.4 | 4576.3 | 26.1 [] |
| Biot-D44 | Biot-PEG-YALGG APPVP SRPGA SPDPF GPPPQ VPSRP NRAPP GVPRT TTSDP-$CONH_2$ | $C_{229}H_{360}N_{64}O_{64}S$ | 5062.7 | 5062.5 | ND |
| Biot-D44-R11A | Biot-PEG-YALGG APPVP SAPGA SPDPF GPPPQ VPSRP NRAPP GVPRI TISDP-$CONH_2$ | | 4823.0 | | [96%] |
| Biot-D54-Dyn2 | Biot-PEG-YAPPI PSRPG PQSVF ANSDL FPAPP QIPSR PVRIP PGIPP GVPSR RPPAA PSRP-$CONH_2$ | | 6025.9 | | [92%] |
| Ac-N-D30 | Ac-YPPVPSRPG ASPDPFGPPP QVPSRPNRAP PG-$CONH_2$ | $C_{147}H_{221}N_{43}O_{40}$ | 3228.7 | 3229.3 | 24.4 [92%] |
| dD15-N | | $C_{249}H_{418}N_{64}O_{73}$ | 5473.1 | 5471.2 | 24.5 [93%] |
| Biot-dD15-N | | $C_{257}H_{430}N_{66}O_{74}S$ | 5657.2 | 5658.4 | 24.9 [92%] |
| dD15-C | | $C_{244}H_{405}N_{65}O_{71}$ | 5382.0 | 5385.1 | 24.8 [85%] |
| Biot-dD15-C | | $C_{266}H_{443}N_{69}O_{77}S$ | 5868.3 | 5868.1 | 24.7 [87%] |

### Supplementary Table 1: Characterization of the synthetized peptides used in this study

**Supplementary Table 2: Proteomics analysis of the proteins interacting with dD15-N.**

| Name | # SH3 domain | BAR domain | Peptide count | Unique peptides | Confidence score | Anova (p) | Max fold change | Description | Accession |
|---|---|---|---|---|---|---|---|---|---|
| **SH3 domain-containing proteins (known Dynamin partners)** | | | | | | | | | |
| SH3kbp1/CIN85 | 3 | NO | 89 | 84 | 264.8 | 1.2E-04 | 207.5 | SH3 domain-containing kinase-binding protein 1 OS=Rattus norvegicus GN=Sh3kbp1 PE=1 SV=2 - [SH3K1_RAT] | Q925Q9;A0A0H2UHD8;M0RBZ7 |
| Intersectin-2 | 5 | NO | 85 | 82 | 245.0 | 2.6E-04 | 118.9 | Intersectin 2 OS=Rattus norvegicus GN=Itsn2 PE=1 SV=2 - [M0R7A6_RAT] | M0R7A6 |
| Intersectin-1 | 5 | NO | 20 | 17 | 52.3 | 1.3E-05 | 60.7 | Intersectin-1 OS=Rattus norvegicus GN=Itsn1 PE=1 SV=3 - [D3ZV52_RAT] | D3ZV52;F1M823;Q9WVE9 |
| Amphiphysin1 | 1 | YES | 20 | 19 | 49.2 | 2.8E-04 | 40.7 | Amphiphysin OS=Rattus norvegicus GN=Amph PE=1 SV=1 - [AMPH_RAT] | O08838;A0A0G2JX32;A0A0G2K5Z4;F1LPP0;Q68FR2 |
| BIN1 (Amph2) | 1 | YES | 18 | 17 | 42.0 | 1.1E-03 | 70.2 | Myc box-dependent-interacting protein 1 OS=Rattus norvegicus GN=Bin1 PE=1 SV=1 - [BIN1_RAT] | O08839;D4A4P1;F1LMX1;Q5HZA7 |
| Endophilin-A1 | 1 | YES | 9 | 6 | 22.7 | 4.7E-03 | 53.5 | Endophilin-A1 OS=Rattus norvegicus GN=Sh3gl2 PE=1 SV=2 - [SH3G2_RAT] | O35179 |
| Endophilin-A2 | 1 | YES | 5 | 2 | 10.1 | 2.8E-03 | 1.7 | Endophilin-A2 OS=Rattus norvegicus GN=Sh3gl1 PE=1 SV=1 - [SH3G1_RAT] | O35964 |
| SNX18 | 1 | YES | 5 | 5 | 10.2 | 2.9E-04 | 16.1 | Sorting nexin OS=Rattus norvegicus GN=Snx18 PE=1 SV=1 - [D3ZZ38_RAT] | D3ZZ38 |
| **SH3 domain-containing proteins (other)** | | | | | | | | | |
| CD2-ap | 3 | NO | 52 | 51 | 125.5 | 8.5E-06 | 124.5 | CD2-associated protein OS=Rattus norvegicus GN=Cd2ap PE=1 SV=2 - [CD2AP_RAT] | F1LRS8 |
| SH3d19 | 5 | NO | 22 | 22 | 55.7 | 1.5E-03 | 42.9 | SH3 domain-containing 19 OS=Rattus norvegicus GN=Sh3d19 PE=1 SV=1 - [D3Z8S0_RAT] | D3Z8S0;D3ZV20 |
| **CCP-related proteins** | | | | | | | | | |
| Eps15l1 | 0 | NO | 92 | 92 | 252.9 | 1.5E-05 | 196.9 | Epidermal growth factor receptor pathway substrate 15-like 1 OS=Rattus norvegicus GN=Eps15l1 PE=1 SV=3 - [D3ZJR1_RAT] | D3ZJR1 |
| Hspa8 | 0 | NO | 23 | 20 | 62.0 | 7.8E-03 | 15.6 | Heat shock cognate 71 kDa protein OS=Rattus norvegicus GN=Hspa8 PE=3 SV=3 - [D4A4S3_RAT] | D4A4S3;A0A0G2JUT0;A0A0G2JVI3;F1LZI1;M0R8M9;M0RCB1;* |
| CHC | 0 | NO | 25 | 25 | 59.5 | 1.3E-04 | 22.7 | Clathrin heavy chain OS=Rattus norvegicus GN=Cltc PE=1 SV=1 - [F1M779_RAT] | F1M779;P11442 |
| AP2a1 | 0 | NO | 24 | 18 | 56.2 | 2.1E-05 | 28.6 | AP-2 complex subunit alpha OS=Rattus norvegicus GN=Ap2a1 PE=1 SV=1 - [D3ZUY8_RAT] | D3ZUY8 |
| AP2a2 | 0 | NO | 22 | 16 | 50.3 | 1.3E-03 | 13.8 | AP-2 complex subunit alpha OS=Rattus norvegicus GN=Ap2a2 PE=1 SV=1 - [Q66HM2_RAT] | Q66HM2;A0A0G2K943;P18484 |
| AP2b1 | 0 | NO | 17 | 17 | 40.9 | 5.4E-07 | 40.0 | AP-2 complex subunit beta OS=Rattus norvegicus GN=Ap2b1 PE=1 SV=1 - [AP2B1_RAT] | P62944;A0A0G2K2V2;G3V9N8;P52303 |
| AP2m1 | 0 | NO | 7 | 7 | 15.4 | 9.4E-04 | 25.5 | AP-2 complex subunit mu OS=Rattus norvegicus GN=Ap2m1 PE=1 SV=1 - [A0A140TAH5_RAT] | A0A140TAH5;P84092 |
| Dnm1 | 0 | NO | 2 | 2 | 4.9 | 1.1E-02 | 27.4 | Dynamin-1 OS=Rattus norvegicus GN=Dnm1 PE=1 SV=2 - [DYN1_RAT] | P21575;A0A0A0MY48;A0A0A0MY49;P39052 |
| **Cytoskeleton proteins** | | | | | | | | | |
| Actg1 | 0 | NO | 13 | 5 | 36.3 | 2.1E-01 | 2.2 | Actin, cytoplasmic 2 OS=Rattus norvegicus GN=Actg1 PE=1 SV=1 - [ACTG_RAT] | P63259;A0A0G2K3K2;P60711 |
| Tubb3 | 0 | NO | 14 | 2 | 30.9 | 3.8E-01 | 1.2 | Tubulin beta-3 chain OS=Rattus norvegicus GN=Tubb3 PE=1 SV=1 - [TBB3_RAT] | Q4QRB4;M0R8B6 |
| Tubb4a | 0 | NO | 13 | 2 | 30.1 | 6.6E-03 | 43.4 | Tubulin beta chain OS=Rattus norvegicus GN=Tubb4a PE=1 SV=1 - [B4F7C2_RAT] | B4F7C2 |
| Tubb5 | 0 | NO | 11 | 2 | 28.9 | 2.5E-03 | 18.0 | Tubulin beta-5 chain OS=Rattus norvegicus GN=Tubb5 PE=1 SV=1 - [TBB5_RAT] | P69897 |
| Tubb2a | 0 | NO | 12 | 3 | 28.3 | 1.6E-03 | 75.4 | Tubulin beta-2A chain OS=Rattus norvegicus GN=Tubb2a PE=1 SV=1 - [TBB2A_RAT] | P85108;Q3KRE8 |
| Tuba1a | 0 | NO | 11 | 4 | 26.4 | 1.5E-02 | 13.3 | Tubulin alpha-1A chain OS=Rattus norvegicus GN=Tuba1a PE=1 SV=1 - [TBA1A_RAT] | P68370;A0A0H2UHM7;F1LUM5;Q68FR8;Q6AY56;Q6AYZ1;Q6P9V9 |
| Actg2 | 0 | NO | 9 | 2 | 22.7 | 3.2E-01 | 2.3 | Actin, gamma-enteric smooth muscle OS=Rattus norvegicus GN=Actg2 PE=2 SV=1 - [ACTH_RAT] | P63269;A0A0G2K4M6;P62738;P68035;P68136 |
| Tuba4a | 0 | NO | 10 | 3 | 20.3 | 1.4E-02 | 14.5 | Tubulin alpha-4A chain OS=Rattus norvegicus GN=Tuba4a PE=1 SV=1 - [TBA4A_RAT] | Q5XIF6 |
| Capza1 | 0 | NO | 3 | 2 | 6.3 | 3.9E-02 | 23.0 | F-actin-capping protein subunit alpha-1 OS=Rattus norvegicus GN=Capza1 PE=1 SV=1 - [CAZA1_RAT] | B2GUZ5 |
| **Other proteins** | | | | | | | | | |
| Calcoco1 | 0 | NO | 16 | 16 | 38.3 | 2.4E-03 | 11.9 | Calcium-binding and coiled-coil domain-containing protein 1 OS=Rattus norvegicus GN=Calcoco1 PE=2 SV=1 - [CACO1_RAT] | Q66HR5;A0A0G2K0S1;A0A1B0GWP3 |
| Hspa5 | 0 | NO | 11 | 8 | 30.1 | 3.8E-02 | 4.1 | 78 kDa glucose-regulated protein OS=Rattus norvegicus GN=Hspa5 PE=1 SV=1 - [GRP78_RAT] | P06761 |
| Rundc3a | 0 | NO | 7 | 7 | 20.0 | 5.9E-04 | 16.0 | RUN domain-containing 3A OS=Rattus norvegicus GN=Rundc3a PE=1 SV=1 - [F1LRZ9_RAT] | F1LRZ9 |
| Hspa9 | 0 | NO | 7 | 6 | 17.2 | 3.2E-04 | 42.1 | Stress-70 protein, mitochondrial OS=Rattus norvegicus GN=Hspa9 PE=1 SV=3 - [GRP75_RAT] | P48721;F1M953 |
| Amy1a | 0 | NO | 6 | 6 | 14.4 | 1.4E-01 | 2.8 | Alpha-amylase OS=Rattus norvegicus GN=Amy1a PE=1 SV=3 - [E9PSQ1_RAT] | E9PSQ1;A0A0G2K6T1;E9PSI7;G3V844;P00689 |
| Atp1a3 | 0 | NO | 3 | 3 | 6.0 | 1.6E-03 | 33.4 | Sodium/potassium-transporting ATPase subunit alpha-3 OS=Rattus norvegicus GN=Atp1a3 PE=1 SV=2 - [AT1A3_RAT] | P06687;P06685;P06686 |
| Ubc | 0 | NO | 2 | 2 | 5.9 | 5.5E-01 | 1.2 | Polyubiquitin-C OS=Rattus norvegicus GN=Ubc PE=1 SV=1 - [UBC_RAT] | Q63429;F1LML2;F1LU69;F1M927;P0CG51;P62982;P62986 |
| Appbp2 | 0 | NO | 3 | 3 | 4.8 | 7.3E-04 | 20.8 | Amyloid protein-binding protein 2 OS=Rattus norvegicus GN=Appbp2 PE=2 SV=1 - [APBP2_RAT] | A5HK05 |

* more accession codes: P0DMW0;P14659;P55063;P63018

**Supplementary Table 3**: p-values for 1-way ANOVA followed by Tukey's multiple comparison tests

1. Tfn-A568 uptake assays (Figure 1E)

| Comparison No treatment vs. | | | Comparison OHT only vs. | | |
|---|---|---|---|---|---|
| OHT only | **** | < 0.0001 | No treatment | **** | < 0.0001 |
| OHT and dyn2-GFP-WT | ns | >0.9999 | OHT and dyn2-GFP-WT | **** | < 0.0001 |
| OHT and dyn2-GFP-ΔPRD | **** | < 0.0001 | OHT and dyn2-GFP-ΔPRD | ns | 0.9893 |
| OHT and dyn2-GFP-ΔCter | **** | < 0.0001 | OHT and dyn2-GFP-ΔCter | ns | 0.2156 |
| OHT and dyn2-GFP-$ABC_{mut}$ | **** | < 0.0001 | OHT and dyn2-GFP-$ABC_{mut}$ | ns | 0.9751 |
| OHT and dyn2-GFP-$A_{mut}$ | ns | 0.9997 | OHT and dyn2-GFP-$A_{mut}$ | **** | < 0.0001 |
| OHT and dyn2-GFP-$B_{mut}$ | **** | < 0.0001 | OHT and dyn2-GFP-$B_{mut}$ | **** | < 0.0001 |
| OHT and dyn2-GFP-$C_{mut}$ | *** | 0.0001 | OHT and dyn2-GFP-$C_{mut}$ | **** | < 0.0001 |
| OHT and dyn2-GFP-$AC_{mut}$ | **** | < 0.0001 | OHT and dyn2-GFP-$AC_{mut}$ | * | 0.0411 |

2. Dyn2-GFP localisation in TKO cells (Figure 1F)

| Comparison dyn2-GFP-WT vs. | | | Comparison dyn2-GFP-ΔPRD vs. | | |
|---|---|---|---|---|---|
| dyn2-GFP-ΔPRD | **** | < 0.0001 | dyn2-GFP-WT | **** | < 0.0001 |
| dyn2-GFP-ΔCter | **** | < 0.0001 | dyn2-GFP-ΔCter | ns | >0.9999 |
| dyn2-GFP-$ABC_{mut}$ | **** | < 0.0001 | dyn2-GFP-$ABC_{mut}$ | ns | >0.9999 |
| dyn2-GFP-$A_{mut}$ | ns | >0.9999 | dyn2-GFP-$A_{mut}$ | **** | < 0.0001 |
| dyn2-GFP-$B_{mut}$ | **** | < 0.0001 | dyn2-GFP-$B_{mut}$ | **** | < 0.0001 |
| dyn2-GFP-$C_{mut}$ | *** | 0.0001 | dyn2-GFP-$C_{mut}$ | **** | < 0.0001 |
| dyn2-GFP-$AC_{mut}$ | **** | < 0.0001 | dyn2-GFP-$AC_{mut}$ | * | 0.0117 |

3. Effect of peptides on endocytic event frequency (Figure 5D)

| Comparison Control WC vs. | | |
|---|---|---|
| No WC recording | ns | 0.5799 |
| GTPγS | ** | 0.0012 |
| D15 NS 1 mM | ns | >0.9999 |
| D15 0.1 mM | ns | 0.9878 |
| D15 1 mM | *** | 0.0002 |
| D30 0.1 mM | * | 0.0338 |
| D44 0.1 mM | * | 0.0359 |
| D44 1 mM | **** | < 0.0001 |
| dD15-N 0.1 mM | * | 0.0476 |
| dD15-N 1 mM | * | 0.0132 |
| dD15-C 0.1 mM | * | 0.0339 |
| dD15-C 1 mM | **** | < 0.0001 |
